# Understanding the bias of compositional microbiome differential abundance estimation

**DOI:** 10.64898/2026.04.28.721392

**Authors:** M.Luz Calle, Meritxell Pujolassos, Antoni Susin

## Abstract

One of the most relevant objectives in microbiome studies is the identification of microbial species that are differentially abundant across conditions. However, the compositional nature of microbiome data complicates this task. Interdependence among components leads to spurious associations when the abundances of each component are analyzed separately. Due to the growing awareness of the challenges of compositional data analysis (CoDA), log-ratio transformations, such as the additive log-ratio (alr) or the centered log-ratio (clr) transformations, have become increasingly popular in microbiome studies.

Several studies have compared the performance of compositional and non-compositional methods through simulations. However, the debate between these two frameworks remains unresolved, creating confusion among researchers. Rather than relying on simulation-based results, this work provides theoretical results that enable a more rigorous and conclusive analysis of the problem, contributing to a better understanding of differential abundance estimation.

We provide theoretical expressions of the bias of differential abundance estimation related to the use of proportions (total sum scaling) and log-ratio transformations (alr and clr) when estimates are interpreted as absolute rather than relative to a reference. The factors that most strongly influence the bias are the magnitude and direction of the effects, the dimension of the composition, the proportion of differentially abundant variables, and the distribution of relative abundances.

The findings of this work strongly support the use of CoDA transformations; however, they also highlight that even when log-ratio transformations are applied, interpreting the results outside of a CoDA framework can still lead to biased conclusions. Among CoDA transformations, alr has several advantages over clr: its reference is more explicit, which reduces the risk of interpreting estimates as absolute rather than relative, and it facilitates the replication of results in independent studies, as it only requires assessing changes relative to the same reference rather than reconstructing the full composition. In this work, we propose a heuristic method for selecting a suitable alr reference component, which will enable a more widespread use of this transformation.

## 1. Introduction

A central goal in the study of the microbiome and its relationship with human health is the identification of microbial species that are differentially abundant between healthy individuals and those with a specific pathology. What might seem like a simple task that could be solved with a statistical test comparing observed abundances in the different groups of individuals is, in fact, one of the main methodological challenges in this field. The primary reason for this difficulty is that microbial abundance data obtained from high-throughput DNA sequencing do not represent the true abundances in the studied ecosystem and can only be interpreted in relative terms (Gloor et al. 2017). Technically, microbial abundances are compositional data constrained to a total (*e*.*g*., the sequencing depth), which is not related to the total abundance in the ecosystem. This constraint on the total causes the observed abundances of each microbial species to be strongly interdependent, such that if one species increases in a sample, the observed relative abundances of the other species will decrease. This interdependence between the different components of the composition leads to spurious associations when the abundances of each component are analyzed separately in relation to a response variable, resulting in a high number of false positives in differential abundance tests. The problem of spurious associations in compositional data was first described by Karl Pearson in 1897 (Pearson 1897), but it was not until 1986 that John Aitchison (Aitchison, 1986) laid the foundations of compositional data analysis (CoDA).

In the context of microbiome research, many authors have emphasized the need to analyze microbial abundance data while accounting for their compositional nature (Gloor and Reid 2016; Gloor et al. 2016; Quinn et al. 2018; Calle 2019; Morton et al. 2019; Greenacre 2021). A wide range of CoDA methods for microbiome analysis are available, such as *ALDEx2* (Fernandes et al. 2014), *ANCOM-BC* (Lin and Peddada 2020a), *fastANCOM* (Zhou C. et al. 2022), *LinDA* (Zhou H. et al. 2022), or *coda4microbiome* (Calle et al. 2023). Other authors, however, advocate for the use of non-compositional methods such as *LEfSe* (Segata et al. 2011), *edgeR* (Robinson et al. 2010), *DESeq2* (Love et al. 2014), *Maslin2* (Mallick et al. 2021) or *metagenomeSeq* (Paulson et al. 2013). The properties of compositional and non-compositional methods have been extensively compared through simulation studies, with results favoring compositional methods in most cases (Lin and Peddada 2020b; Nearing et al. 2020; Yang and Chen 2022; Weiss et al. 2017), whereas in others their validity and reliability are called into question (Wirbel et al. 2024; Gamboa-Tuz et al. 2025; Pelto et al. 2025). The controversy between these two approaches is substantial, causing considerable confusion among researchers who wish to analyze the data of their microbiome studies with the most reliable methods available.

The bias introduced when estimating differential abundance from relative abundances has been widely described in the literature (e.g., Morton et al. 2019; Lin and Peddada 2020a). This has popularized the use of the centered log-ratio (clr) transformation in the microbiome field, but often without proper consideration of its implications, leading to incorrect interpretations of the estimates as absolute rather than relative to the reference defined by this transformation.

In this work, we compare the performance of non-compositional estimates based on relative abundances (total sum scaling) with log-ratio transformations, namely the additive log-ratio (alr) and the centered log-ratio (clr). We first show that the estimation of differential abundance is not identifiable unless a reference is specified, either explicitly or implicitly. In the case of log-ratio transformations, the reference is explicit: the geometric mean of the effects in the case of clr, and one of the components in the case of alr. Although the estimates from total sum scaling (TSS) are often assumed to be absolute, this normalization also implicitly specifies a reference: the ratio of total abundances between the two environments. The estimates from any of these approaches must be interpreted as variations with respect to the reference. If this is ignored and the results are interpreted as absolute rather than relative, all three approaches yield biased estimates.

The questions we address in this article are, first, which transformation provides estimates closer to the true effects, and second, which reference is more meaningful and facilitates the interpretation of the results.

Rather than basing our work on simulations, in this study we undertake a theoretical examination of these methods that enable a more rigorous and conclusive analysis of the problem. Specifically, we present the theoretical expressions for the bias in differential abundance estimation associated with the use of proportions (total sum scaling) and log-ratio transformations (alr and clr) when estimates are interpreted as absolute rather than relative to a reference. We describe the conditions required for each transformation to give unbiased estimates and in which situations the bias may have a greater impact and in which other ones it may be negligible. The factors that most strongly influence the bias are the magnitude and direction of the effects, the dimension of the composition, the proportion of differentially abundant variables, and the distribution of relative abundances. In the case of the alr transformation, it is well known that the condition for obtaining unbiased results is to use a reference component that is not associated with the response variable.

The findings of this work strongly support the use of CoDA transformations, but they also highlight that even when log-ratio transformations are applied, interpreting the results outside of a CoDA framework can still lead to biased conclusions. The study of differential abundance requires a fully compositional view of both the problem and its results, recognizing that the parameters of interest (fold changes) are only estimable up to a scale factor that cannot be identified from the data. Due to this identifiability problem, differential abundance estimation is not well-defined without the specification of a reference. Conclusions will depend on the reference choice and results obtained under different references will not be directly comparable.

Among CoDA transformations, alr has several advantages over clr, but its use has been hindered by the challenge of selecting the reference component. In this work we propose a heuristic method to obtain a suitable alr reference and explore its performance through simulations. The algorithm is implemented as an R function of the *coda4microbiome* package available at CRAN (https://cran.r-project.org/web/packages/coda4microbiome/).

## 4. Methods

In this section, we begin by describing the estimators of log fold change based on three different transformations, TSS, CLR and ALR, and derive their expected values and bias (Subsection 2.1). Then we show that the TSS bias is larger than the CLR bias under certain conditions (Subsection 2.2), and that fold changes are only estimable up to a scaling factor (Subsection 2.3). In Subsection 2.4, we propose an algorithm to obtain a suitable alr reference. Finally, Subsections 2.5 and 2.6 describe the different scenarios used in the analytical and simulation studies.

### 2.1. Bias of differential abundance estimates

We consider a composition of *D* microbial species that are present in two different environments A and B. The true absolute abundances in environment *X* ∈ {*A, B*} are denoted by 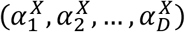, where 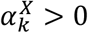 for each *k* ∈ {1, …, *D*}. The true relative abundances in *X* are denoted by 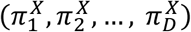, where 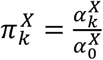 and 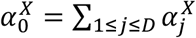 is the total abundance in environment *X*. By definition, 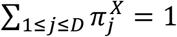.

We are interested in estimating the differences in microbial abundances between both environments, and, more precisely, the fold change effects, denoted by ***F*** = (*F*_1_, *F*_2_, …, *F*_*D*_), and defined as 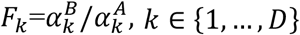.

Given ***F***, the true absolute abundances in B can be expressed as 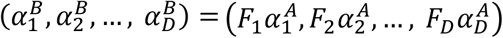. Moreover, the true relative abundances in B are given by 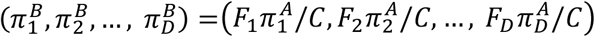, where 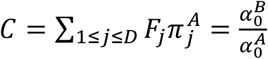 is the ratio between total abundances in environments B and A. Figure 1 illustrates the notation with an example.

**Fig. 1.**
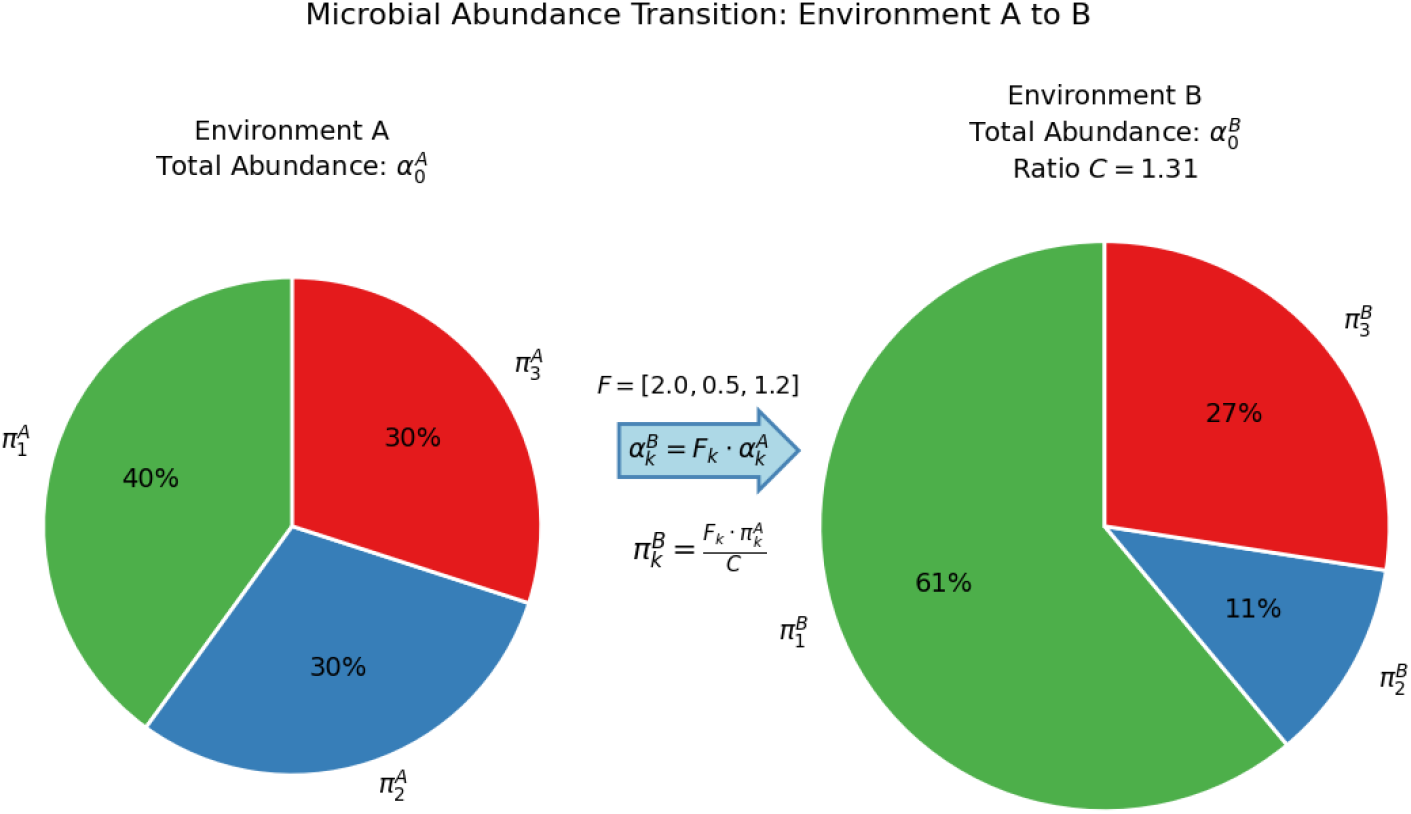
Example and schematic representation of absolute and relative microbial abundances across two environments. Environment A (left) features relative abundances 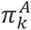 with the total area of the pie chart corresponding to the total absolute abundance 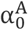. Environment B (right) illustrates the altered relative abundances 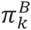 after applying the species-specific fold change effects *F*_*k*_. The scaled size of the pie chart in Environment B reflects the total abundance ratio 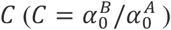, visually demonstrating the mathematical relationship 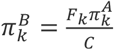

The goal of univariate differential abundance testing is to identify those microbial species with significantly different abundance in environments A and B. This corresponds to testing the null hypothesis *F*_*k*_ = 1, or, equivalently, log(*F*_*k*_) = 0, for each component *k* ∈ {1, …, *D*}. Estimation of fold change values, *F*_*k*_, requires sampling from both environments.

We denote by 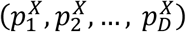 the *D-*dimensional random vector of observed relative abundances, 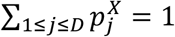, in environment *X* ∈ {*A, B*} and assume the sampling process was completely at random, *i*.*e*. 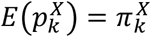, for *k* ∈ {1, …, *D*}.

Below we detail three different approaches, TSS, CLR and ALR, for estimating the parameter of interest, log(*F*_*k*_), for differential abundance testing. For each approach, we provide the expression of the expected value of the estimator and their bias:

#### 2.1.1 TSS estimation

Normalizing by the total number of counts per sample to obtain relative abundances in a composition is also known as “total sum scaling”. The log fold change for each component *k* can be estimated as the difference between the sample means of the log relative abundances in both environments. We denote this estimator as *logF*_*k TSS*_:

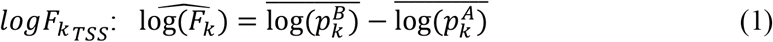

##### Expected value of *logF*_***kTSS***_

###### Proposition 1.

The expected value of the estimator of the log fold change, log(*F*_*k*_), using the TSS as defined in Equation 1 is given by:

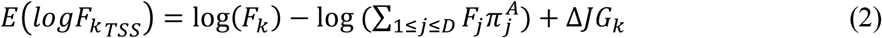

where 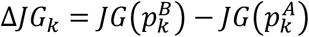 is the difference between Jensen’s gaps defined as *JG*(*p*_*k*_) = *E*(log(*p*_*k*_)) − log(*E*(*p*_*k*_)) (Jensen 1906).

*Proof:*

The expected value of *logF*_*kTSS*_ is given by:

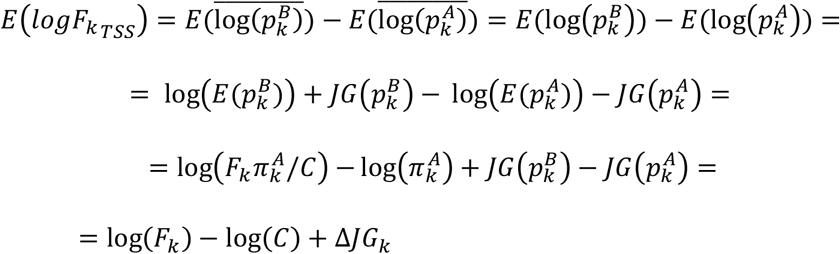

with 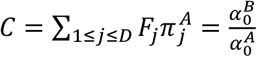 .

##### Bias of *logF*_*kTSS*_

According to proposition 1 and assuming that the difference between Jensen’s gaps is negligible (Supplementary material), the bias of the TSS estimation is:

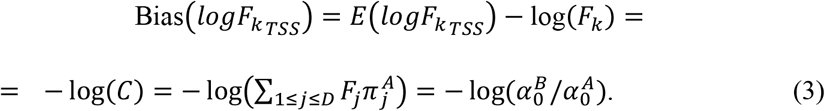

Alternatively, it can be seen as providing an unbiased estimator of log(*F*_*k*_/*C*), where *C* is the total abundance ratio between both environments.

#### 2.1.2 CLR estimation

We consider the centered log-ratio (clr) transformation of a composition. Given a vector of observed relative abundances, ***p*** = (*p*_1_, *p*_2_, …, *p*_*D*_), with ∑*p*_*k*_ = 1, the clr transformation is defined as:

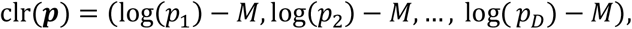

where 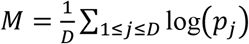

The log fold change for each component *k* can be estimated as the difference between the sample means of the clr-transformed relative abundances in both environments. We denote this estimation as *logF*_*kCLR*_:

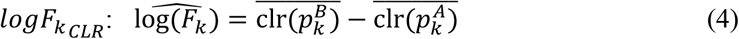

##### Expected value of *logF*_***kCLR***_

###### Proposition 2.

The expected value of the clr estimator of the log fold change for component *k* is given by:

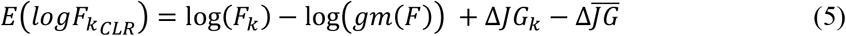

where *gm*(*F*) denotes the geometric mean of ***F*** and 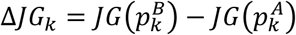 is the difference between Jensen’s gaps for component *k* and 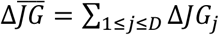 is the mean of Jensen’s gaps difference for all *D* components.

*Proof:*

To derive the expected value of *logF*_*kCLR*_ we need first to obtain the expression of the expected value of *M*:

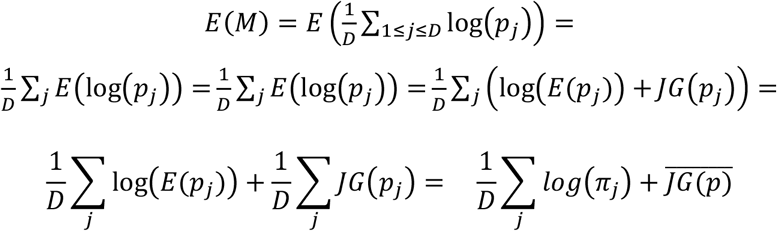

Again, we can assume that 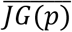 is negligible, and thus, 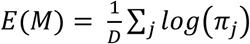, the geometric mean of the true relative abundances in ***π***.

The expected value of the clr estimator of the log fold change for component *k* is given by:

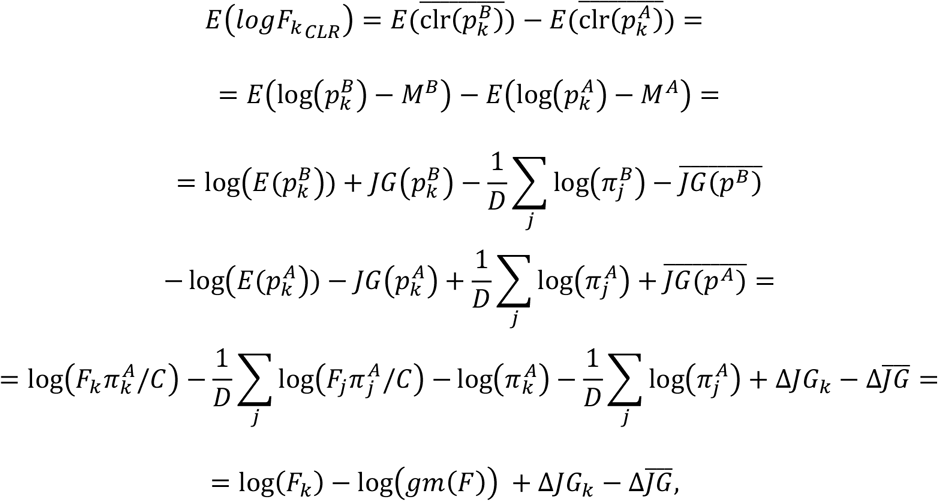

where *gm*(*F*) denotes the geometric mean of *F* and 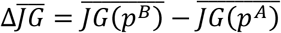.

##### Bias of *logF*_***kCLR***_

Assuming again that the difference between Jensen’s gaps is negligible, the bias of the CLR estimation is:

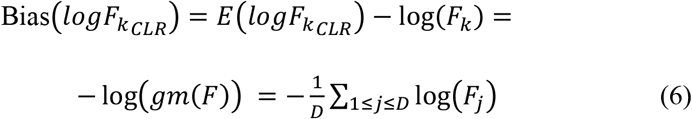

Alternatively, it can be seen as providing an unbiased estimator of log(*F*_*j*_/*gm*(*F*)), where *gm*(*F*) is the geometric mean of effects.

#### 2.1.3 ALR estimation

Finally, we consider the additive log-ratio (alr) transformation of a composition that consists of taking one of the components as the reference for the others. Given a vector of observed relative abundances, ***p*** = (*p*_1_, *p*_2_, …, *p*_*D*_), with ∑*p*_*k*_ = 1, the alr transformation with the last component, *D*, as the reference is defined as:

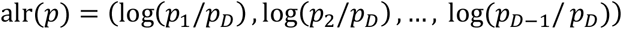

The log fold change for each component *k* can be estimated as the difference between the sample means of the alr-transformed relative abundances in both environments.

We denote this estimation as *logF*_*kALR*_:

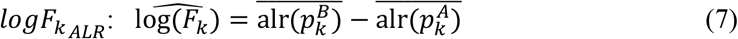

##### Expected value of *logF*_***kALR***_

To be more general, instead of considering the last component as the reference, we will denote by *Ref* the index of the reference component.

###### Proposition 3.

The expected value of the alr estimator of the log fold change for component *k* taking component *Ref* as the reference is given by:

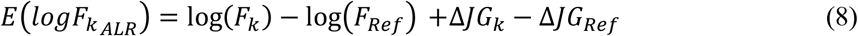

*Proof:*

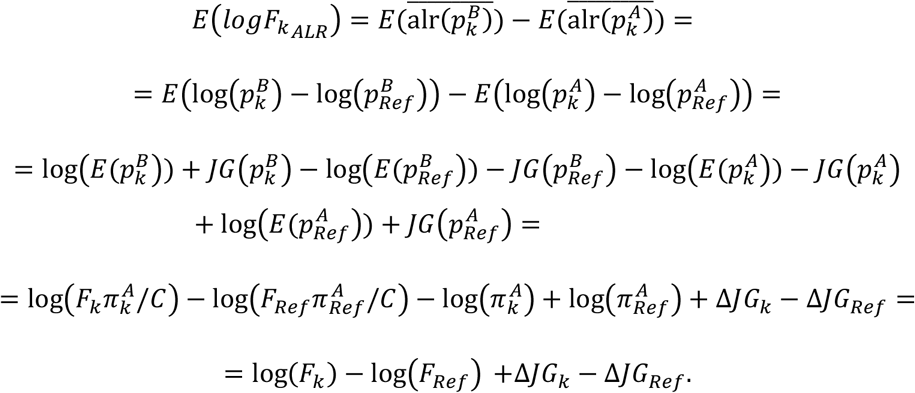

##### Bias of *logF*_***kALR***_

Assuming again that the difference between Jensen’s gaps is negligible, the bias of the alr estimation is given by minus the log fold change of the reference variable:

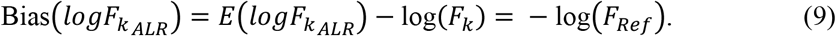

Alternatively, it can be seen as providing an unbiased estimator of log(*F*_*j*_/*F*_*Ref*_), where *F*_*Ref*_ is the effect of the reference component.

### 2.2. Comparison between the TSS and CLR biases

While the CLR bias (equation 6) only depends on the values of the fold changes, the TSS bias (equation 3) depends on both the fold changes and the distribution of true relative abundances in environment A.

#### Proposition 4.

If the true relative abundances in environment A are uniformly distributed, *i*.*e*. 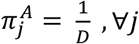, the absolute bias of TSS estimation is larger than the absolute bias of the CLR estimation:

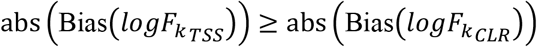

*Proof:*

Because of Jensen’s inequality, the following holds:

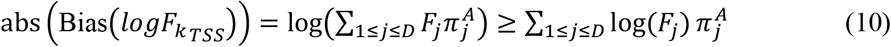

Still, the inequality, that will prove that the absolute value of the TSS bias is larger than the absolute value of the CLR bias does not always hold:

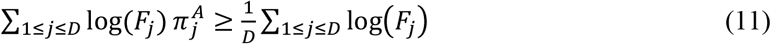

### 2.3. Scaling factor

Let’s consider a situation where all components are affected by a global effect, *S, e*.*g*. a reduction of the total microbial load due to the use of antibiotics. In this case, the fold change of each component can be expressed as the global effect, *S*, multiplied by the specific effect of each component, denoted by *F*_*k*_:

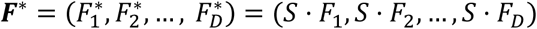

According to equations (1, 4 and 6), the expected values of the three estimation methods, not including the Jensen’s gap terms, can be rewritten as follows:

Expected value of the estimator of log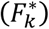 using TSS estimation:

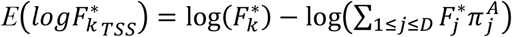

Since 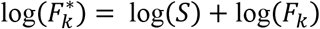, the term log(*S*) cancels and thus,

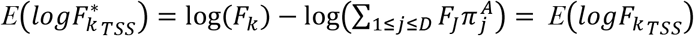

Similarly, for the CLR estimation:

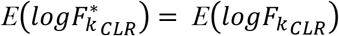

and also, for the ALR estimation:

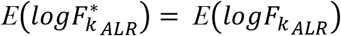

In all cases, the global effect term *S* cancels out. We will discuss the implications of these expressions in the Results section.

### 2.4. Heuristic method for obtaining a good reference for ALR estimation

Based on the bias formulas (Equations 3, 6, and 9), the method with the simplest bias, and therefore the easiest to control, is presumably the ALR method. For the ALR bias to be null, it is necessary to select a reference component that is not associated with the response variable, in our case, one that has a similar abundance in both environments, *i*.*e*. with a fold change equal to 1, so that log(*F*_*Ref*_) = 0.

Under the assumption that most of the components have a null effect, *i*.*e*. log(*F*_*k*_) = 0, for most *k*, we propose the following algorithm to find a suitable reference component that results in an unbiased alr estimation. In the results section we will discuss the situation where all components are affected by a scaling factor and thus, all of them are differentially abundant between the two environments.

The algorithm relies on the density distribution of the estimated log fold change of all *D* parts of the composition, *i*.*e*., the log fold change of each microbial species abundance, using the TSS or the CLR estimations. We first describe the algorithm using the TSS estimation:

Since the bias of the TSS estimation is constant for all parts of the composition (Equation 3), the distribution of TSS estimates, *logF*_*kTSS*_, 1 ≤ *k* ≤ *D*, is identical to the distribution of true fold changes, log(*F*_*k*_), except for a shift factor equal to the bias −log(*C*), where 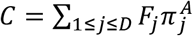.

Under the assumption that most of the components have a null effect, the density distribution of true fold changes, log(*F*_*k*_), should have a mode equal to zero. Because of the shift factor, the distribution of estimated effects, *logF*_*kTSS*_, will have a mode different from zero. However, the components with values near this mode correspond to those with values close to zero in the true fold change distribution and are therefore unlikely to be differentially abundant.

We propose to select as the reference component for ALR estimation, the variable, denoted by *Ref*, with an estimated effect close to the mode of the *logF*_*kTSS*_ distribution. By default, the algorithm considers the 10 percent of components closest to the mode and selects the one with the lowest variance. The algorithm is implemented in the R function RefALR within the *coda4microbiome* package (Calle et al. 2024).

The same reasoning and procedure can be applied using clr estimation and the *logFk*_*CLR*_ distribution.

### 2.5. Analytical study

Given the expression of the bias for the TSS, CLR and ALR estimation methods, we conducted a study to explore their behavior across different parameter settings. This analysis did not require empirical sample data, as exact formulas are available to compute the bias.

According to the bias formulas, the relevant parameters to be considered are the dimension of the composition, *D*; the number or percentage of differentially abundant (DA) variables or components, *K*_*DA*_ and %*K*_*DA*_; the vector of true relative abundances in environment A, denoted by *π* ^*A*^; and the vector of fold changes, *F*.

For each scenario, we considered compositions with different dimension *D* ∈ {10, 50, 100, 500, 1000}. The first *K*_*DA*_ components were those differentially abundant between the two environments, where *K*_*DA*_ depends on the dimension *D* of the composition. We considered %*K*_*DA*_ ∈ {10%, 20%, 30%, 40%}.

For the true relative abundances, ***π***^***A***^, we considered two scenarios:

(1) a uniform distribution with constant values 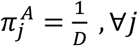;

(2) a non-uniform skewed distribution where 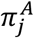 is sampled from (exp(*λ* = 1))^2^.

We considered two different ways of assigning the fold change to the first *K*_*DA*_ components (the rest of components have a fold change equal to 1):

a. Constant: the same fold change value for all DA variables:

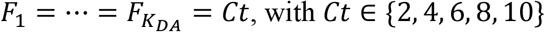
b. Uniform sampling:

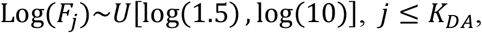

The direction of the change (increase or decrease) of each differential abundant component was stablished by multiplying the log fold changes by a sign *s* = 1_{_*U*_[_0,1_]_<*p*_}_ − 1_{_*U*_[_0,1_]_>*p*_}_, where *p* is the proportion of positive log fold changes, with *p* ∈ {0.5, 0.7, 1}.

### 2.7. Simulation study

We performed a simulation study to evaluate the mean squared error of the three transformations and to explore the ability of the proposed algorithm, RefALR, to identify a suitable reference variable for the alr transformation. Using the same parameters as in the previous bias analysis, we simulated 100 samples from each environment, A and B, drawing from a Dirichlet distribution with parameter ***α***^***X***^ = ***π***^***X***^***ϕ***, for *X* ∈ {*A, B*}. The precision parameter was set to ***ϕ*** = *D*/2 to introduce overdispersion, thereby generating more realistic abundance data.

To avoid the issue of zeros in the logarithmic transformation, and for the sake of simplicity and computational efficiency in the simulations, a small constant value (10^−8^) was added to all relative abundances in the taxa table. Components with more than 80% zeros were removed.

## 3. Results

### 3.1. Identifiability of effects and differential abundance testing

In the Methods section, we showed that, regardless of the transformation used –TSS, CLR or ALR – the parameter of interest, the fold change vector (*F*_1_, *F*_2_, …, *F*_*D*_), is identifiable only up to a multiplicative constant. Equivalently, the log fold change vector (log(*F*_1_), log(*F*_2_), …, log(*F*_*D*_)) cannot be distinguished from a translation by a constant, (log(*F*_1_) + *K*, log(*F*_2_) + *K*, …, log(*F*_*D*_) + *K*). Due to this identifiability problem, the differential abundance hypothesis of interest, H_0_: log(*F*_*j*_) = 0, are not well-defined and require the specification of a reference. The conclusions of the test will depend on the reference choice, and results obtained under different references will not be directly comparable. This is exemplified in Figure 2, where the log fold change vector in (b) is obtained from (a) by applying a shift to the reference value. When using the reference in (a), components 5 and 6 are not differentially abundant; however, under the reference in (b), they appear to be the most differentially abundant.

**Fig. 2.**
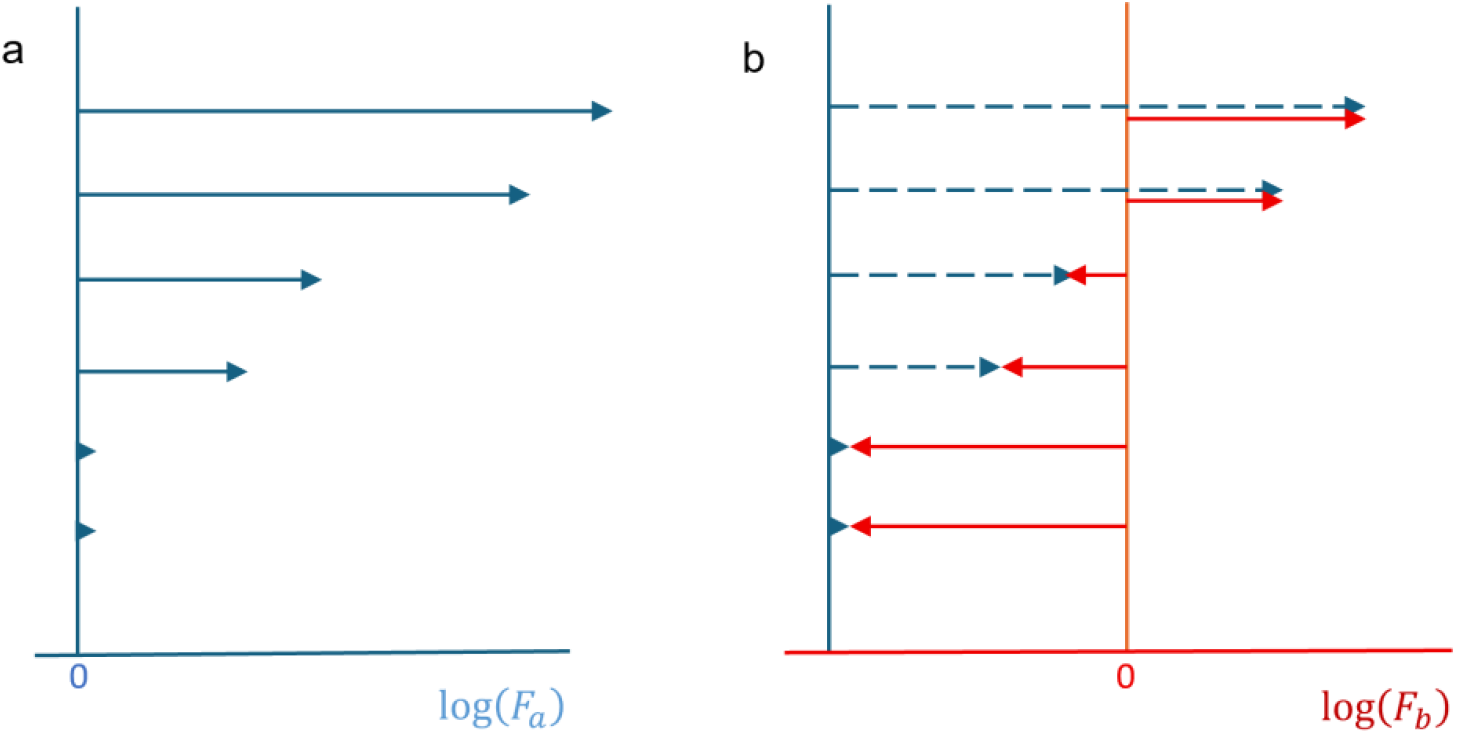
Example of a composition with 6 components. In (a), log(*F*_*a*_) = (10, 9,5,4,0,0) and in (b), log(*F*_*b*_) = (4, 3, −1, −2, −6, −6). The difference between (a) and (b) is a shift equal to −6. In this example, if the CLR was used, the shift would be equal to −log(*gm*(*F*)) = −4.66; and it would be equal to −log(*C*) = −log (∑_1≤*j*≤*D*_ *F*_*j*_*π*_*j*_) = −8.52 for the TSS transformation, assuming *π*_*j*_ = 1/6.

CoDA approaches acknowledge the identifiability issue, explicitly define a reference, and interpret the results with respect to that reference. When the CLR transformation is used, the reference is the geometric mean of effects, the estimated effects are expressed in relation to this reference and the hypothesis tested is H_0_: log(*F*_*j*_/*gm*(*F*)) = 0. Under the ALR transformation, the hypothesis tested is H_0_: log(*F*_*j*_/*F*_*Ref*_) = 0.

It is commonly but incorrectly assumed that non-CoDA transformations neither define nor require a reference and directly test the hypothesis of interest, H_0_: log(*F*_*j*_) = 0; however, this is not correct. The TSS transformation implicitly uses the total abundance change as its reference, and the hypothesis addressed in this case is H_0_: log(*F*_*j*_/*C*) = 0.

Given the different reference adopted by each of the three methods, discrepancies in the results are expected, as they correspond to different hypothesis tests and are therefore not expected to coincide. This is usually ignored in empirical and simulation studies that compare the performance of the different approaches.

In the following sections, the discussion adopts a non-CoDA interpretation of results, as it is common in differential abundance testing, regardless of whether log-ratio transformations (CLR and ALR) are used or only total-sum normalization (TSS). This aims to illustrate the challenges of interpreting results ignoring the implicit or explicit reference of each method.

### 3.2. Behavior of the bias of the TSS, CLR, and ALR methods

In this section, we discuss the main differences between the three estimation methods, TSS, CLR and ALR, based on the expressions of their bias to estimate differential abundance (equations 3, 6 and 9). From now on, when we refer to bias, we will mean the bias in absolute value, regardless of its direction. We compare the magnitude of the bias of the three approaches in different scenarios and identify those situations in which the bias is negligible.

The first aspect to highlight is the simplicity of the bias of the ALR method, which depends only on the log fold change of the reference variable. In contrast, the bias of the CLR method depends on the effect of all the variables in the composition, specifically, on the arithmetic mean of the log fold changes. Finally, the bias of the TSS method depends on both, the effect (fold change) of all the variables in the composition and on the distribution of the true relative abundances in environment A.

#### 3.2.1. Conditions for unbiasedness

These are the conditions that need to be verified for each method to be an unbiased estimate of fold changes:

TSS: Same total abundances of environments A and B, *i*.*e*. 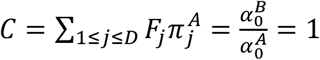.

CLR: The arithmetic mean of log fold changes is zero, *i*.*e*. 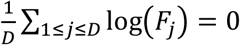.

ALR: The reference variable is not differentially abundant between the two environments, *i*.*e. F*_*Ref*_ = 1 or, equivalently, log(*F*_*Ref*_) = 0.

Though none of these conditions can be verified from the observed data, it is possible to anticipate that some of these conditions may be more difficult to justify than others. In any case, their validity relies on the assumption of an additional condition that could be reasonable or not within the context of the specific phenomenon being studied. For example, for the CLR method, in specific contexts it might be reasonable to assume that some variables increase in abundance while others decrease, resulting in a nearly unbiased estimate.

In the case of the ALR, if the hypothesis that most of the variables are not differentially abundant is reasonable, it is possible to find a suitable reference variable that yields unbiased results (proposed algorithm, Subsection 2.4).

The most stringent condition for unbiasedness is that of the TSS estimation, as it requires that the abundances in both environments are identical, a condition difficult to justify since it depends on both the magnitude of fold changes and the distribution of abundances.

#### 3.2.2. Evaluation of the bias of the TSS and CLR methods

We evaluated the bias formulas of the TSS and CLR methods across different parameter settings. In this analysis, we did not consider the ALR method since its bias only depends on the log fold change of the reference variable. The results are summarized in Figures 3 and 4. They show the mean absolute bias for different percentages of significant variables (panel columns), and for different percentages of significant variables with an increase in abundance, *i*.*e*. fold change > 1 (panel rows). In Figure 3, the x-axis of each plot corresponds to the dimension of the composition *D* and the fold change for each significant component was randomly sampled. In Figure 4, all significant components had the same fold change *F* (x-axis of each plot), and the dimension of the composition was fixed to *D* = 100. Figure 3a and Figure 4a correspond to a uniform distribution of relative abundances and Figure 3b and Figure 4b to a non-uniform distribution.

**Fig. 3.**
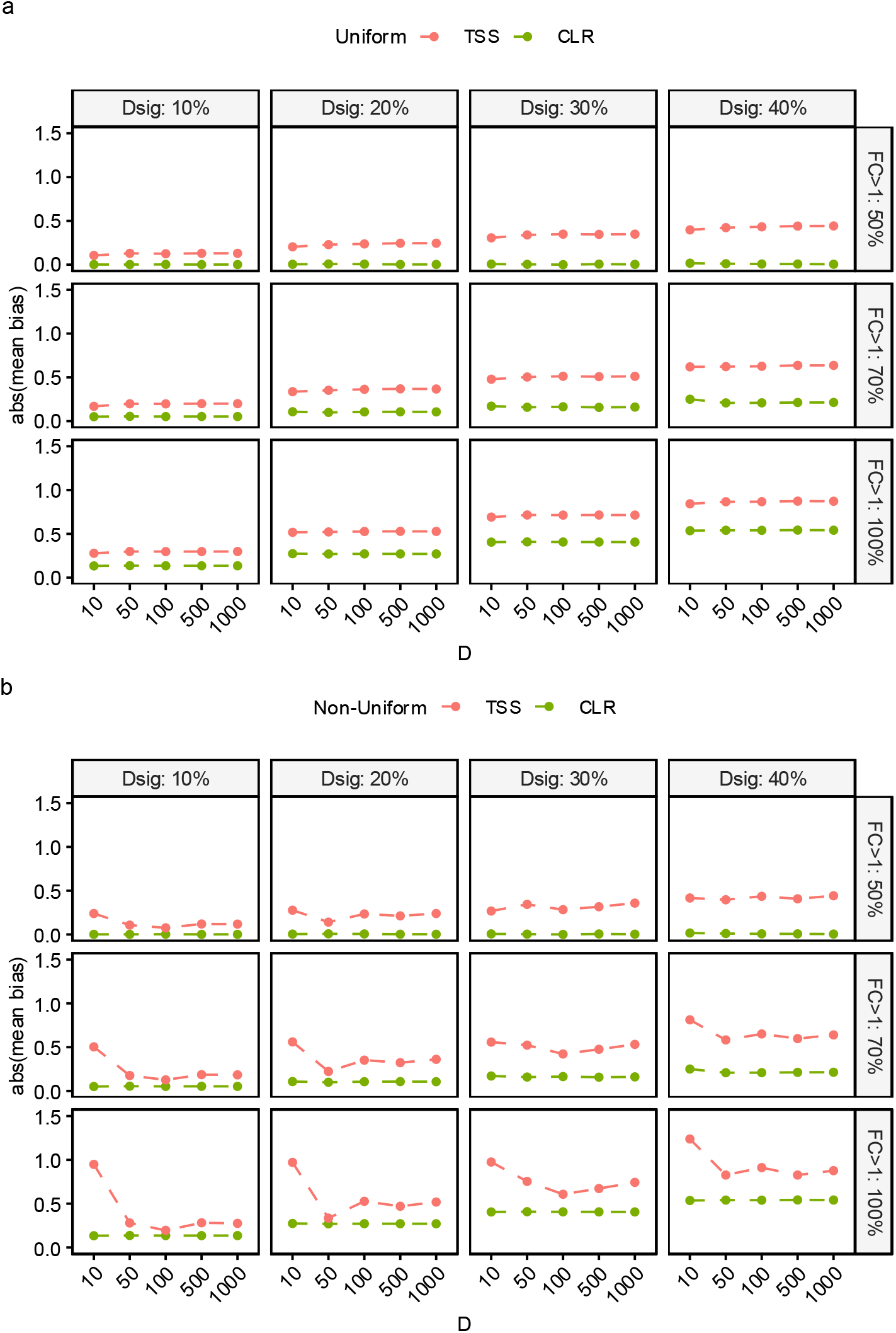
Theoretical mean bias (in absolute terms) of for TSS (red) and CLR (green) methods under uniform (a) and non-uniform (b) distribution of abundances. The x-axis of each plot corresponds to the dimension of the composition (D), the panel columns correspond to the proportion of differentially abundant variables (10%, 20% or 40%, Dsig); and the panel rows correspond to the direction of the effects (50%, 70% or 100% of significant variables with FC>1)

**Fig. 4.**
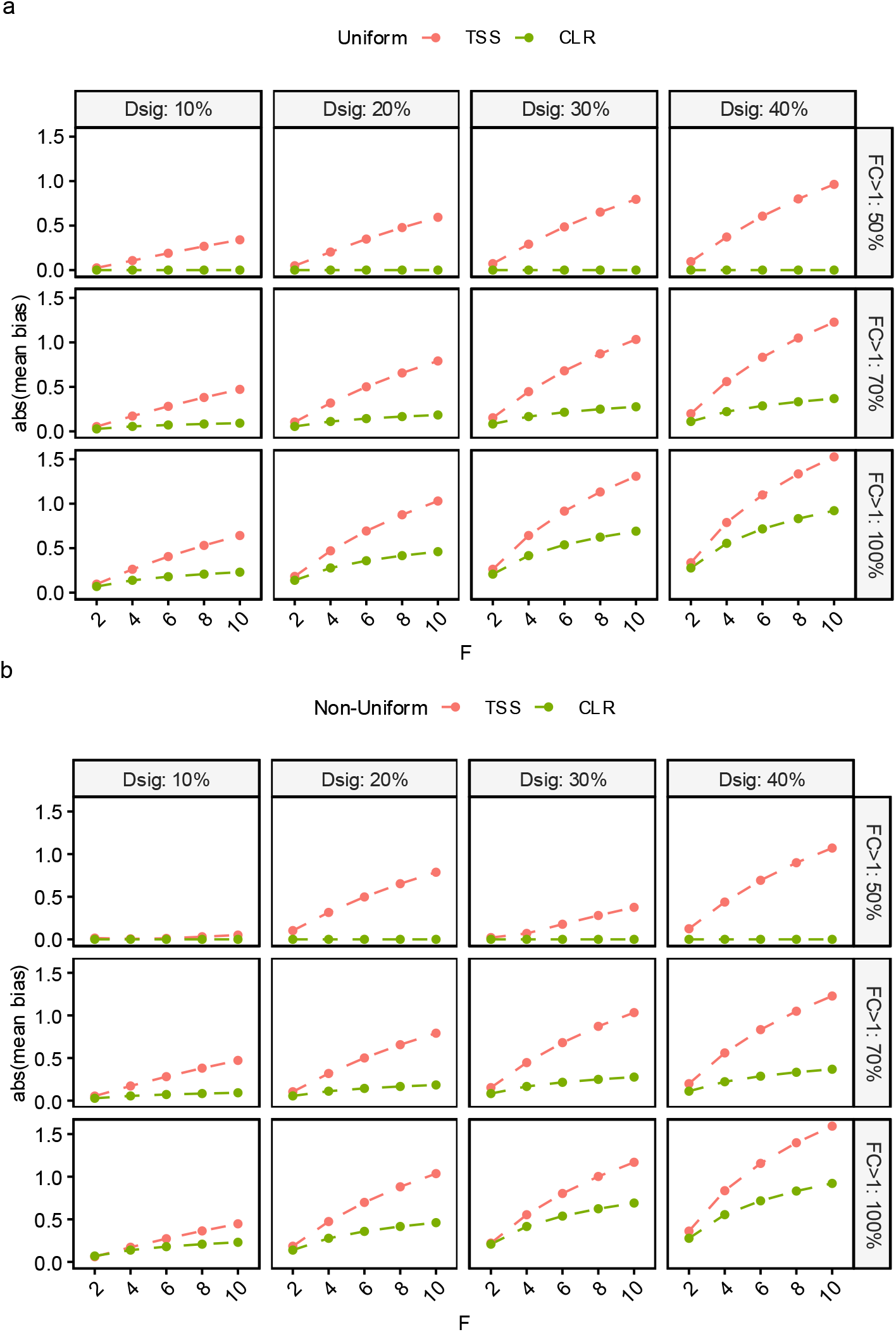
Theoretical mean bias (in absolute terms) of for TSS (red) and CLR (green) methods under uniform (a) and non-uniform (b) distribution of abundances. The x-axis of each plot corresponds to the fold change of each significant component (F), the panel columns correspond to the proportion of differentially abundant variables (10%, 20% or 40%, Dsig); and the panel rows correspond to the direction of the effects (50%, 70% or 100% of significant variables with FC>1)

Below we discuss the most relevant results for each parameter:

##### Distribution of true relative abundances

As demonstrated in *Proposition 4* (Methods Section), if the abundance distribution is uniform, the bias of the TSS method will always be greater than that of the CLR method. Figures 3a and 4a confirm this theoretical result for the different scenarios considered. Although it cannot be proven theoretically, this is likely to happen also when the abundance distribution is not uniform. We confirmed this experimentally in the evaluation of the biases under a non-uniform distribution of relative abundances, and in all cases the bias of the TSS method was greater than that of the CLR method (Figures 3b and 4b).

##### Dimension of the composition and proportion of differentially abundant variables

For the same proportion of differentially abundant variables, %*K*_*DA*_, the bias remains quite stable across the different values of *D*, especially for the CLR bias, which is almost constant, while for the TSS method, the bias is larger for smaller values of *D* when the distribution of relative abundances is non-uniform (Figure 3b).

##### Direction of the effects

For both the TSS and CLR methods, the smallest bias is observed when the proportions of significant variables with positive log fold changes (increasing abundance) and negative log fold changes (decreasing abundance) are similar (Figures 3 and 4, first row of the panel). In this case, the CLR bias is practically zero as was to be expected, since the bias of the CLR method corresponds to the mean of the log fold changes.

The bias of both methods increases as the proportion of variables changing in the same direction increases. The largest bias occurs when all significant variables change in the same direction (Figures 3 and 4, last row of the panel).

##### Magnitude of fold changes

In the case of the TSS method, the bias increases with the magnitude of the fold change (assuming that all significant variables have the same fold change). For the CLR method, this is also true except in cases where there is a similar proportion of variables that increase and decrease, since, as we mentioned earlier, in this situation the bias is practically zero regardless of the composition size (Figure 4).

### 3.3. Selection of a suitable reference variable

The simulation results, summarized in Figure 5, indicate that the proposed algorithm for selecting the reference variable for the ALR transformation performs very well. The plot shows the proportion of correct identifications by the RefALR algorithm, where correctness is defined as selecting a reference component unrelated to the response variable, *i*.*e*. same abundance in both environments. For the uniform distribution of abundances, the success rate is nearly 100% in all scenarios, with an average success rate of 99%. The performance for the non-uniform distribution is also very good, with an average success rate of 98%. The only case with a success rate below 90% (bottom-right corner) corresponds to a small number of components (D=10), a large proportion of significant variables (Dsig=40%) and all them varying in the same direction (100% FC>1).

**Fig. 5.**
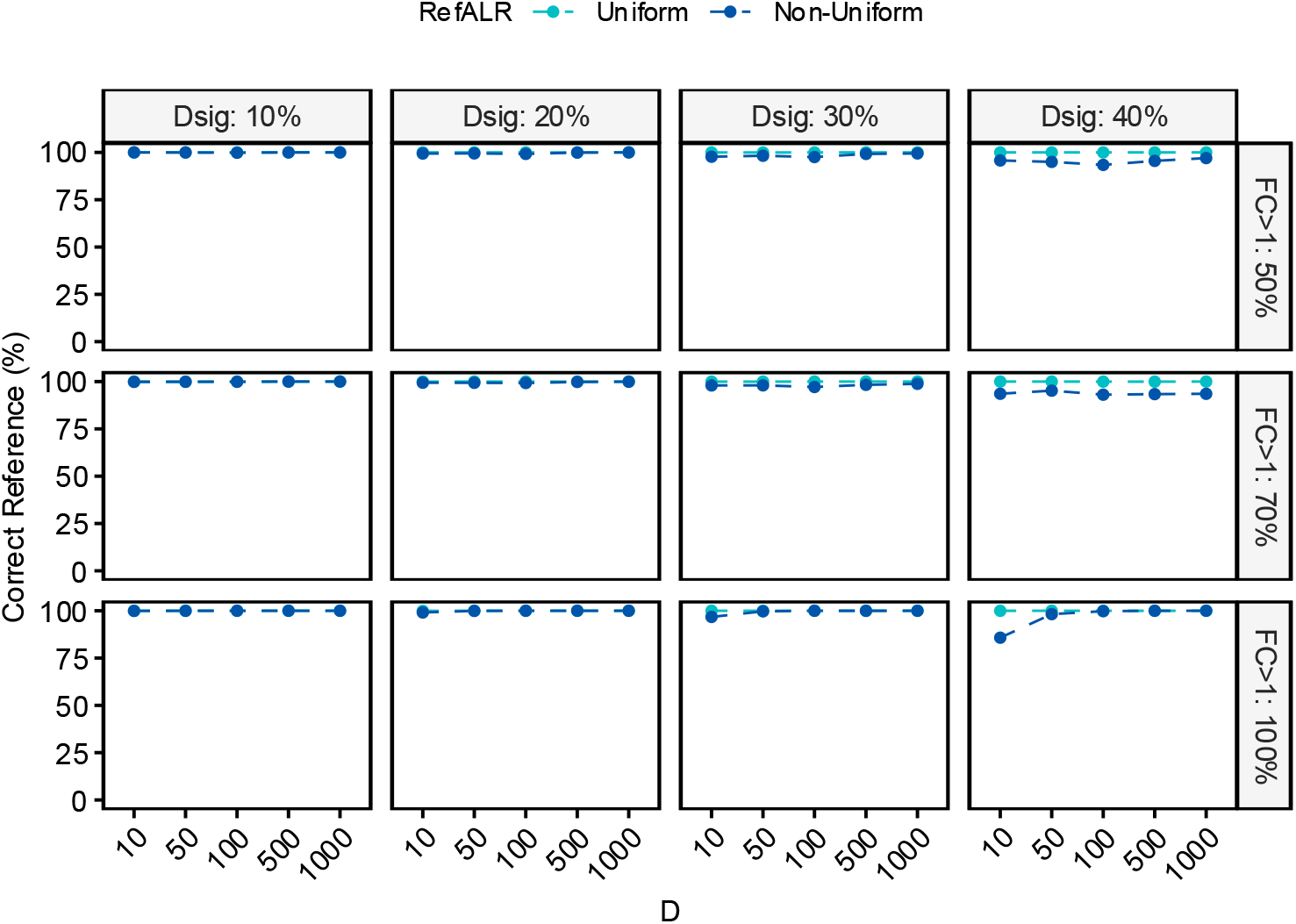
Success rate of the RefALR algorithm, where success is defined as the selection of a reference variable not associated with the response variable, under uniform (light blue) and non-uniform (dark blue) distribution of abundances. The x-axis of each plot corresponds to the dimension of the composition (D), the panel columns correspond to the proportion of differentially abundant variables (10%, 20% or 40%, Dsig); and the panel rows correspond to the direction of the effects (50%, 70% or 100% of significant variables with FC>1)

### 3.4. Mean Squared Error

The quality of an estimator is determined not only by its bias but also by its variance. An estimator with low bias produces values that are, on average, close to the true parameter, but if its variance is high, those estimates can fluctuate widely from sample to sample, making it unreliable in practice. The mean squared error, defined as MSE = Bias^2^ + Variance, combines both aspects and provides a global measure of the accuracy of an estimator. Below we provide the MSE results for the three data transformation methods.

Figure 6 compares the MSE between TSS and CLR estimation for (6a) uniform and (6b) non-uniform distribution of relative abundances and random fold changes. In all scenarios, the TSS MSE is considerably larger than that of the CLR method, with larger MSE for smaller compositions. As for the CLR method, in the case of a uniform distribution (Figure 6a), the MSE is indeed very small across all scenarios. This is because, as we saw previously, the bias is practically negligible and, in addition, the variability associated with CLR normalization is very low, since it is computed using the arithmetic mean of the log fold changes. In the case of a non-uniform distribution (Figure 6b), the MSE associated with the CLR method is slightly higher. This may be because, in this scenario, a considerable proportion of components have a very low mean abundance, which leads to increased variability. In any case, the MSE of the CLR method remains lower than that of the TSS method.

**Fig. 6.**
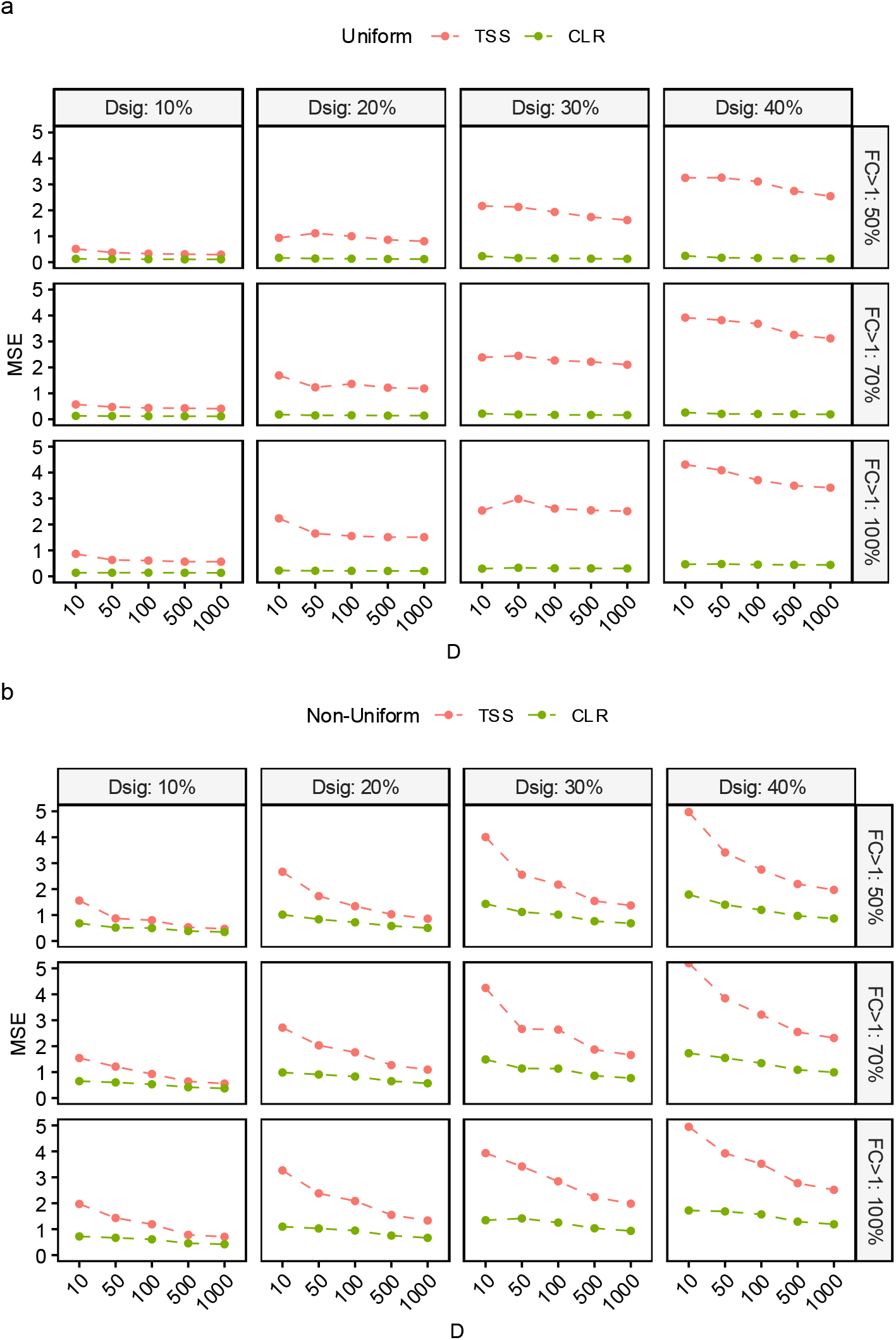
Mean squared error (MSE) for TSS (red) and CLR (green) methods under uniform (a) and non-uniform (b) distribution of abundances. The x-axis of each plot corresponds to the dimension of the composition (D), the panel columns correspond to the proportion of differentially abundant variables (10%, 20% or 40%, Dsig); and the panel rows correspond to the direction of the effects (50%, 70% or 100% of significant variables with FC>1)

In Figure 7 we compare the MSE of the CLR and the ALR using as reference the candidate obtained using the proposed algorithm, RefALR, implemented as a function in R package *coda4microbiome* (Calle et al. 2024).

**Fig. 7.**
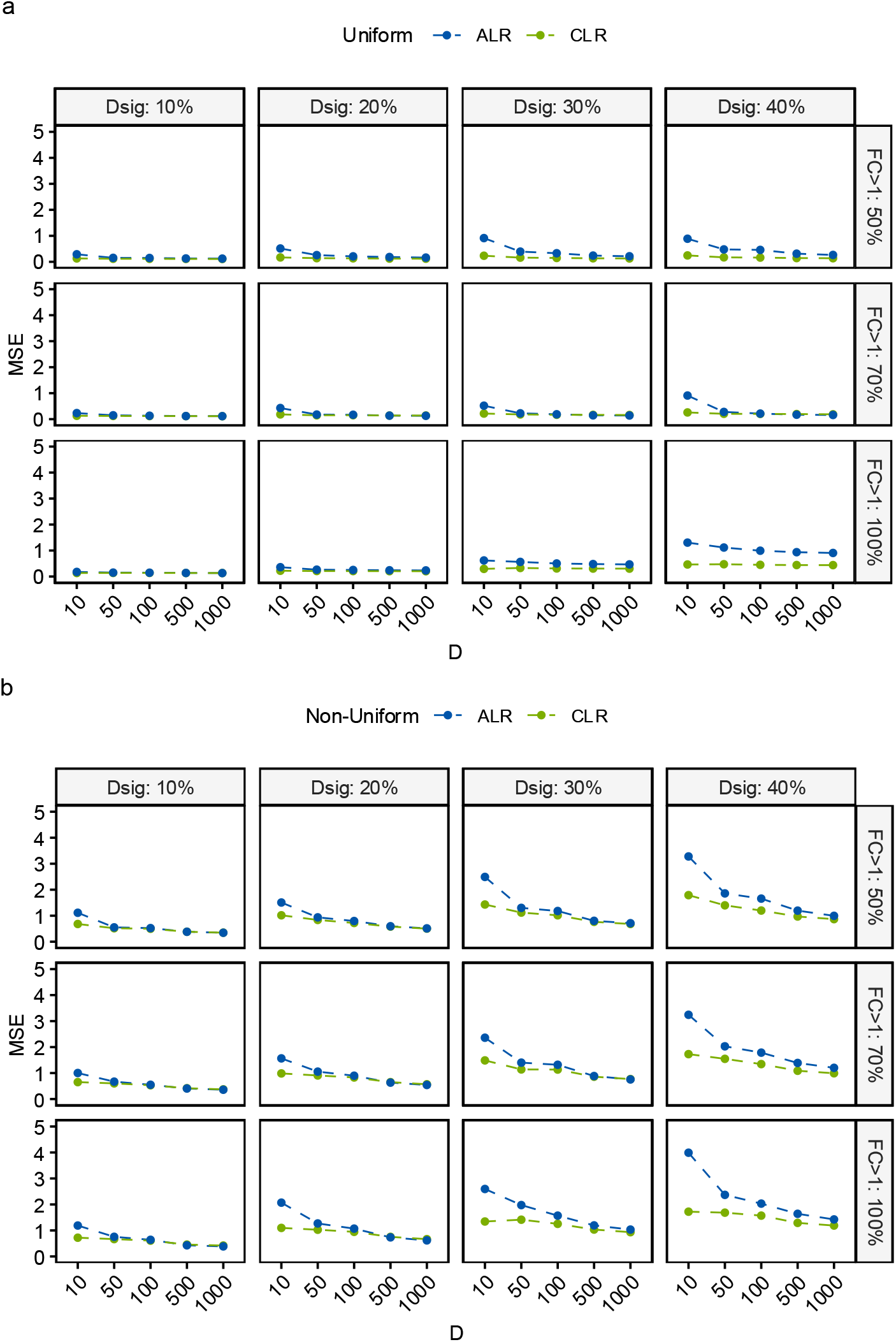
Mean squared error (MSE) for ALR (blue) and CLR (green) methods under uniform (a) and non-uniform (b) distribution of abundances. The x-axis of each plot corresponds to the dimension of the composition (D), the panel columns correspond to the proportion of differentially abundant variables (10%, 20% or 40%, Dsig); and the panel rows correspond to the direction of the effects (50%, 70% or 100% of significant variables with FC>1)

For abundances uniformly distributed, both methods have very good performance, *i*.*e*. small MSE, in all scenarios. Their performance is essentially identical, except for the case with a large percentage of significant variables (Dsig = 40%) and all of them changing in the same direction (100% FC>1), where ALR has slightly larger MSE than CLR.

For the case of non-uniform distribution of abundances, the performance is again very similar between both methods, except when the dimension of the composition is small (D=10) and the percentage of significant variables is moderate (Dsig >= 30%). In these cases, the CLR performs better than the ALR. This could be explained by the higher number of zeros in the abundance matrix, leading to poorly informative variables that may introduce variability in both estimation methods, but with a greater impact on the ALR method.

## 4. Discussion

In this work, we have addressed one of the most relevant and complex topics in microbiome research: quantifying the increase or decrease in the abundance of a microorganism between two experimental conditions, for example, between patients suffering from a disease and healthy individuals. This quantification enables the identification of specific bacteria that are differentially abundant in a disease state or following an intervention, such as a diet or treatment.

The main difficulty in this analysis lies in the fact that abundance data obtained from DNA sequencing are compositional and therefore contain relative, rather than absolute, information about the abundance of species in each sample. As discussed, this implies that the magnitude of the differences in abundance (fold change) cannot be quantified in absolute terms and can only be determined up to a multiplicative factor. Therefore, regardless of the transformation applied to the data, the results must be interpreted with this limitation in mind.

Several studies have compared, through simulation analyses, the performance of differential abundance tests under different data transformations. However, the validity of simulation-based results may be questioned in relation to their design and the scenarios considered. In contrast, this work provides theoretical results that allow for a more rigorous and conclusive analysis of the problem.

Our results show that the CLR and ALR transformations perform substantially better than the simple TSS transformation. We demonstrate that, if the distribution of relative abundances is uniform, the bias introduced by the TSS transformation in estimating fold change is always greater than the bias associated with CLR. In the case of a non-uniform distribution of relative abundances, there is no general theoretical relationship between the two biases; however, in all the scenarios analyzed, the bias of the TSS transformation was greater than that of CLR, which in many cases is null or negligible.

Specifically, we observed that when, between the two experimental conditions, there is a similar proportion of species that increase in abundance and others that decrease, the bias after the CLR transformation is practically zero. Bias in CLR only appears when most components vary in the same direction (either increasing or decreasing). This bias is not problematic if CLR-based estimates are interpreted as increases or decreases relative to the average variation across all components. For instance, a negative log fold change does not necessarily imply that abundance has decreased, but rather that it has changed (either increased or decreased) less than the average change across all components.

In contrast, for TSS it is difficult to find a simple interpretation that avoids potential misunderstandings due to bias, since the reference center used by TSS does not have a clear meaning.

The ALR transformation is, in principle, the optimal transformation, since if the reference component is chosen appropriately, the fold change estimate is unbiased. In practice, however, this transformation suffers from two difficulties: the choice of a suitable component, namely, one that is independent of the response variable, and the variability introduced by normalization with respect to this reference component. In this work, we have proposed a heuristic method to obtain a suitable reference component for the ALR transformation that, under the assumption that most of the components have a null effect, yields an unbiased estimate of the log fold change. We have verified the very good performance of this algorithm in cases where the proportion of differentially abundant components is moderate. Since this condition cannot be verified from the data, the results of the ALR transformation must be interpreted with caution, assuming that the fold change estimates are relative to the changes of the chosen reference component.

Considering the MSE, which encompasses both the bias and the variability of the estimates, we have found that the CLR transformation shows the best performance, closely followed by the ALR transformation using the algorithm proposed in this article for selecting the reference component. The results show that the use of the TSS transformation is not advisable in this context, given the high MSE values obtained.

The identifiability problem of log fold change effects, which are only identifiable up to an additive constant, prevents any possibility of correcting the bias using only relative abundance data; this would require experimental information on total abundances. Some authors advocate the use of quantitative microbiome profiling, which includes experimental procedures to estimate microbial load, to help overcome the compositional limitations of sequencing data (Stämmler et al. 2016, Tkacz et al. 2018, Lloréns-Rico et al. 2021, Nishijima et al. 2025). However, technical sources of variability can introduce substantial additional bias related to the quantification method which limits the practical utility of this approach (Galazzo et al. 2020).

With this study, we aim to contribute to a better understanding of the problem of estimating differential abundance between two environments by providing theoretical expressions of the bias of the main data transformations and introducing a new algorithm to obtain a reference variable for the ALR transformation. The theoretical findings we present strongly support the use of CoDA transformations, but they also highlight that there is no easy solution to this complex problem and that, in any case, the results must be interpreted relative to a given reference. In fact, the estimators are biased relative to the true effects, but they are unbiased if the reference is made explicit.

Among CoDA approaches, the CLR transformation is straightforward. However, in practice, many researchers interpret its results as absolute effects rather than changes relative to the mean, which can lead to incorrect conclusions. In addition, results are harder to replicate, since replication requires reconstructing the full composition rather than evaluating only the variables of interest, which may be costly or infeasible.

In contrast, ALR is simple and keeps the reference variable explicit. Replication in independent studies only requires assessing changes relative to the same reference, without reconstructing the full composition. We have proposed an algorithm for selecting the reference variable in the ALR transformation, which will enable a more widespread use of this transformation. When external evidence supports that most components have no effect, the algorithm provides a reference component that is independent of the outcome, and in this case ALR estimates may be interpreted as approximations of true effects. Otherwise, these estimates should be interpreted as variations relative to the reference variable.

We have focused on addressing the bias in the estimation of effects introduced by the analysis method and, in particular, by the data transformation. It is important to note that microbiome studies may involve other sources of experimental bias that we have not addressed (McLaren et al. 2019, Hu et al. 2022, Maghini et al. 2023).

This work was motivated by a specific problem in the microbiome field; however, the results presented are general and apply to any field involving compositional data where differential abundance testing is required.

## Supporting information

Supplemental proof proposition 1

## Data availability

All data supporting the findings of this study are available within the paper and its Supplementary Information.

