## Supplemental proof proposition 1 for "Understanding the bias of compositional microbiome differential abundance estimation"

**Calle ML, Pujolassos M and Susin A**

### Suppl information 1.

$\Delta JG_k = JG(p_k^B) - JG(p_k^A)$ *is approximately equal to zero*

$$\Delta JG_k = JG(p_k^B) - JG(p_k^A) =$$

$$E(\log(p_k^B)) - \log\left(E(p_k^B)\right) - E(\log(p_k^A)) + \log\left(E(p_k^A)\right) \cong (*)$$

$$E(\log(F_k p_k^A)) - \log\left(E(F_k p_k^A)\right) - E(\log(p_k^A)) + \log\left(E(p_k^A)\right) =$$

$$\log(F_k) + E(\log(p_k^A)) - \log(F_k) - \log\left(E(p_k^A)\right) - E(\log(p_k^A)) + \log\left(E(p_k^A)\right) = 0$$

(*) Since $\pi_k^B = F_k \pi_k^A$, the empirical relative abundandes: $p_k^B \cong F_k p_k^A$
